# Lung tissue viscoelasticity is preserved with bleomycin-induced fibrosis in mice

**DOI:** 10.1101/2025.09.06.674583

**Authors:** Leilani R. Astrab, Riley T. Hannan, Mackenzie L. Skelton, Jeffrey M. Sturek, Steven R. Caliari

**Author notes:** Co-corresponding authors (S.R. Caliari), (J.M. Sturek).

## Abstract

In pulmonary fibrosis, excessive scar tissue accumulates in the alveolar interstitial space, impairing gas exchange and compromising lung function. This fibrotic remodeling results in tissue stiffening, but more complex lung mechanical properties critical to tissue function, such as viscoelasticity and stress relaxation, remain poorly defined. To address this gap, we use the bleomycin aged mouse model to characterize both bulk and spatially-resolved viscoelastic mechanical properties of normal and fibrotic lungs. Our analysis reveals that while bleomycin-induced fibrosis leads to heterogeneously increased lung stiffness, viscoelasticity as measured by tan delta (ratio of loss to storage modulus) and stress relaxation timescales remains remarkably consistent as a function of both age and bleomycin treatment. This unexpected preservation of viscoelasticity despite fibrotic stiffening highlights a previously underappreciated mechanical phenotype of fibrotic lungs. To model these distinct mechanical features *in vitro*, we utilize a hyaluronic acid-based hydrogel system that largely recapitulates the viscoelastic mechanical properties observed in both normal and fibrotic lungs. These findings provide new insight into the mechanical consequences of fibrosis and establish a tunable *in vitro* hydrogel platform mimicking key tissue viscoelastic properties.

## 1. INTRODUCTION

Idiopathic pulmonary fibrosis (IPF) is the most common of interstitial lung diseases (ILDs), characterized by chronic and irreversible formation of scar tissue with no known etiology [1]. While the direct cause of IPF is unknown, the sustained activation of extracellular matrix (ECM)-producing myofibroblasts [2,3] results in an excess and disorganized accumulation of ECM constituents like collagen and hyaluronic acid (HA). As scar tissue accumulates in the alveolar interstitial space, lung tissue becomes progressively stiffer and less deformable, compromising its ability to withstand cyclic stretch. This mechanical dysfunction reduces lung capacity, impairs oxygen diffusion, and ultimately contributes to progressive respiratory decline and failure [4]. Understanding the interplay between ECM remodeling, tissue mechanics, and fibrosis progression is crucial for developing targeted therapies to mitigate disease progression and preserve lung function.

In recent decades, our understanding of fibrosis pathology has been expanded by the adaptation of *in vitro* biomaterial-based models that better mimic mechanical changes occurring in disease. Biomaterial models have facilitated the study of biomimetic mechanical dosing on cells in a manner not possible when cells are cultured on supraphysiologically-stiff glass or plastic. These studies have revealed the role of mechanical stiffening as not just a byproduct of fibrosis progression, but also a driver in the disease [5–7]. Further, there is a growing appreciation that stiffness alone does not fully capture the complexity of the mechanical environment cells experience *in vivo*. In particular, viscoelasticity, the time-dependent mechanical behavior of materials that exhibit both solid-like and liquid-like properties, has emerged as a critical regulator of cellular behavior [8–13]. In some cases, changes in viscoelasticity have been shown to outweigh stiffness in directing cellular activity [10]. Viscoelastic properties can be quantified using storage modulus (G’), which demonstrates the solid-like component of materials (energy stored), and loss modulus (G”) which represents the liquid-like behavior of materials (energy dissipated). Tissues, as well as biological materials from reconstituted ECM, typically display a ratio of loss to storage moduli (G”/G’ = tan delta) of at least 0.1 [12]. Furthermore, viscoelastic materials are stress-relaxing, meaning they dissipate applied stress over time in response to constant strain, which has been shown to influence cell dynamics [13]. Thus, it is of utmost importance to develop *in vitro* models that accurately depict the full repertoire of mechanical changes that occur in fibrosis to better understand the role they play in disease.

Animal models have been invaluable for providing benchmarking design criteria for these *in vitro* systems, and enabling insights into how the cellular and mechanical environment of lung tissue evolves with fibrotic disease progression. Among these, the bleomycin mouse model is the most widely used approach for investigating IPF *in vivo* [14]. Bleomycin-treated mice have been shown to develop several key features of human pulmonary fibrosis, including immune cell infiltration, activation of myofibroblasts, increased collagen production, and aberrant ECM remodeling [15,16]. However, a primary challenge in using this model to investigate IPF is that it spontaneously resolves over time, failing to capture the chronic and progressive nature of the disease [17–19]. To address this issue, studies have shown that aged mice demonstrate persistent scar tissue formation, as well as more severe fibrosis [20,21]. These findings highlight the aged mouse bleomycin model as a more pathophysiologically relevant tool for studying human IPF, making it better suited for investigating the impact of lung tissue changes in fibrosis.

While the bleomycin model has been extensively used to study pulmonary fibrosis at the cellular and molecular levels, dynamic changes in lung tissue mechanics remain largely unexplored. To date, most studies have focused on documenting increases in tissue stiffness, whereas changes in viscoelasticity—critical for proper lung function—have been largely overlooked. While one study identified regions of mature fibrosis in decellularized human lungs to be more linearly elastic than healthy areas or fibroblastic foci, this measurement doesn’t directly translate to tissue viscoelasticity nor does it inform changes to bulk lung mechanics [22]. Given that viscoelasticity influences how lung tissue dissipates mechanical stress and regulates cell–matrix interactions, characterizing its alterations during fibrosis is essential. Such insights could ultimately guide the development of biomaterial models that more accurately capture the mechanical environment of fibrotic lung tissue.

Here, we employ an aged mouse model to investigate how bleomycin-induced fibrosis influences lung tissue stiffness and viscoelasticity. To comprehensively assess how fibrotic remodeling impacts lung mechanics, we employed complementary techniques: rheology to evaluate bulk viscoelastic behavior and nanoindentation to capture spatial heterogeneity at the microscale. This multi-scale approach enables us to determine whether fibrosis alters not only the overall ability of the lung to dissipate energy and recover from deformation, but also how these properties vary across tissue regions. Finally, we adapt an HA-based hydrogel system previously developed in our lab [8,9,23] to model the lung mechanics quantified in normal and bleomycin-treated mice. These findings provide new insights into the mechanical consequences of fibrosis and inform biomaterial design for generating disease-relevant *in vitro* models.

## 2. MATERIALS AND METHODS

### 2.1 Animal care and bleomycin model

All animal studies were approved by the University of Virginia Animal Care and Use committee. C57BL/6J male mice were purchased from Jackson Laboratory at 8-10 weeks old and aged to 15 months before bleomycin instillation. Mice receiving bleomycin were sedated and bleomycin (0.3 U/kg) was administered trans-orally similar to previously described methods [24]. Briefly, after anesthesia, mice were suspended by the upper incisors and their tongues retracted before 50 µL of bleomycin solution was pipetted to the posterior oropharynx. While maintaining tongue retraction, the nares were covered until the bleomycin solution was aspirated into the lungs.

After bleomycin administration, mice were weighed daily and clinically scored based off body weight, appearance, and activity [25]. Body weight was scored on a scale of 0 to 3, with 0 indicating no weight loss, 1 indicating < 10% weight loss, 2 corresponding to 10–19% weight loss, and 3 representing > 20% weight loss. Appearance was scored based off grooming and eye condition on a scale from 0 to 2, with 0 indicating normal appearance, 1 corresponding to a ruffled or ungroomed coat, and 2 indicating sunken, closed eyes. Activity was evaluated on a scale of 0 to 3, 0 being normal activity, 1 indicating decreased activity, 2 meaning inactive and less alert, and 3 indicating very restless. Humane euthanasia endpoint was set as 20% weight loss, a total score of 8, or a maximum score in two categories. Mice were euthanized 3 weeks post-bleomycin instillation and lungs harvested for subsequent mechanical testing and histological evaluation.

### 2.2 Rheological characterization of lung tissue

Following harvest, lungs were kept in phosphate-buffered saline (PBS) on ice until mechanical characterization (∼ 1 hr). Bulk lung mechanics were determined with a stress-controlled DHR-2 rheometer (TA Instruments), equipped with a sandblasted 8 mm diameter parallel plate geometry at 25 °C. Left lung lobe mechanical properties were evaluated using an oscillatory time sweep (1 Hz, 1% strain), oscillatory frequency sweep (0.1–10 Hz, 1% strain), and stress relaxation test (5% strain, 3 min).

### 2.3 Nanoindentation characterization of lung tissue

After rheological characterization, the same left lung lobes were used to evaluate spatial heterogeneity via nanoindentation. To mechanically characterize the inside of the lung, lobes were aligned in a rat heart slicer from Zivic Instruments with 0.5 mm spaces (model HSRS005-1) and cut in half along the coronal plane. Lobes were attached to 15 mm diameter petri dishes using a silicone-based waterproof adhesive and allowed to adhere for 5 min before adding PBS to submerge lungs. Indentation tests were performed on an Optics11 Life Piuma nanoindenter using a 50 μm borosilicate glass probe attached to a cantilever with a spring constant of 0.47 N/m.

Indentations were made to a depth of 4 μm using a constant time ramp of 2 sec. E’ and E” were determined using dynamic mechanical analysis (DMA) at a frequency of 1 Hz across an 8 x 8 matrix with 200 µm spacing between points. Three points in different lung areas were selected for frequency sweeps (0.1-10 Hz) and stress relaxation measurements (10 μm depth, 3 min). Lungs were then removed from plates and processed for histology.

### 2.4 Histology and immunofluorescent staining

Following mechanical characterization, left lung lobes were fixed in 4% paraformaldehyde for 20 min then submerged in 30% sucrose until no longer buoyant (minimum 24 hr). Lungs were then cryosectioned and underwent hematoxylin and eosin (H&E) and Masson’s trichrome staining.

Ashcroft scoring was performed double-blinded by two trained, independent researchers using Masson’s Trichrome-stained slides. Each mouse was scored using 10 random fields from a whole-slide tile scan. The Ashcroft scoring system is an established method for fibrosis disease-staging in bleomycin-treated mice [26–28]. This system consists of eight grades of fibrosis, with a score of 0 for healthy lung. Scoring is performed on 1 mm^2^ (∼ 20x magnification) fields of lung parenchyma, with airway and larger blood vessel areas excluded heuristically by pathologists. Grades 1 – 4 are characterized by interstitial tissue thickening without compromised alveolar architecture, while grades 5 – 8 are characterized by presence of fibrotic masses and progress in severity with the proportion of parenchyma replaced by interstitial tissue, up to complete loss of normal tissue at a score of 8.

Cryosections were dehydrated in PBS, permeabilized in PBS + 0.3% Triton-X100, blocked in PBS + 5% bovine serum albumin + 5% normal goat serum, blocked with excess avidin and then biotin, and stained for HABP-Biotin (1:100) and rabbit anti-Col1a1 (1:200) in blocking buffer for 2.5 hr at 25°C. Secondaries of steptavidin-AlexaFluor647 (1:500), goat anti-rabbit AlexaFluor555 (1:1000) as well as conjugated anti-aSMA-AlexaFluor488 (1:100) were added in PBS + 5% BSA overnight at 4°C. Finally, sections were counterstained with Hoechst 33342 at a final concentration of 0.004 mg/mL in PBS for 5 min at room temperature. Slides were mounted and imaged on a Leica THUNDER TIRF. Images were acquired using an HC PL FLUOTAR L 20x/0,40 CORR PH1 objective and Leica K8 sCMOS camera using appropriate excitation and emission filters to distinguish Hoechst, AlexaFluor488, AlexaFluor555, and AlexaFluor647.

### 2.5 Norbornene-modified hyaluronic acid (NorHA) synthesis

NorHA was prepared via 4-(4,6-dimethoxy-1,3,5-triazin-2-yl)-4-methylmorpholinium chloride (DMTMM)-mediated amide coupling between sodium hyaluronate (Lifecore, 19 kDa) and 5-norbornene-2-methylamine [29]. HA (0.6 mmol) was dissolved in a solution of 5-norbornene-2-methylamine (0.2 mmol) and ultrapure water and the pH adjusted to 6 using HCl. DMTMM (0.3 mmol) was then added to the solution, which was subsequently allowed to react at 40°C overnight. The reaction’s pH was adjusted to 4 using HCl then promptly precipitated in cold ethanol and readjusted to pH 7 with NaOH. The precipitate was then redissolved in water and dialyzed against brine for 1 day, then water for 1 day, and lyophilized. The degree of modification was 19% as determined by ^1^H NMR (Bruker Neo 400 MHz NMR Spectrometer) (**Fig. S1**).

### 2.6 β-CD-HDA synthesis

β-cyclodextrin (β-CD) hexamethylene diamine (β-CD-HDA) was synthesized using a previously outlined method [30]. Briefly, *p*-toluenesulfonyl chloride (TosCl) dissolved in acetonitrile was added dropwise to a solution of β-CD (5:4 molar ratio of TosCl:CD) at 25°C and allowed to react for 2 hr. The reaction was then cooled on ice and an aqueous NaOH solution added dropwise (3.1:1 molar ratio of NaOH to CD). The reaction proceeded for 30 min at 25°C followed by the addition of ammonium chloride to reach a pH of 8.5. The solution was cooled on ice, precipitated using cold water and acetone, and dried overnight. The CD-Tos product was then charged with hexamethylene diamine (HDA) (4 g/g CD-Tos) and dimethylformamide (DMF) (5 mL/g CD-Tos), then reacted under nitrogen at 80°C for 12 hr. The reaction solution was then precipitated in cold acetone (5 × 50 mL acetone/1 g CD-Tos), washed with cold diethyl ether (3 × 100 mL), and dried. The β-CD-HDA product was confirmed using ^1^H NMR (**Fig. S2**).

### 2.7 Cyclodextrin-modified hyaluronic acid (CDHA) synthesis

β-CDHA was synthesized via benzotriazole-1-yloxytris-(dimethylamino)phosphonium hexafluorophosphate (BOP)-mediated coupling of β-CD-HDA with HA tetrabutyl ammonium salt (HA-TBA). The reaction was allowed to proceed for 3 hr at 25°C in anhydrous dimethyl sulfoxide (DMSO), then quenched with cold water. The reaction solution was then dialyzed (molecular weight cutoff 6–8 kDa) for 5 days, filtered, dialyzed an additional 5 days, frozen, and lyophilized. The degree of modification was determined to be 25% by ^1^H NMR (**Fig. S3**).

### 2.8 HA hydrogel fabrication and mechanical characterization

Hydrogels were fabricated according to previously described methods [8,9,23,31]. Briefly, thiolated adamantane peptide (Ad-KKKCG) and dithiothreitol (DTT) crosslinker were reduced using tris(2-carboxyethyl)phosphine (TCEP) (0.5:1 mol TCEP:thiol) prior to hydrogel precursor preparation. Concentrated stock solutions of CDHA and the thiolated adamantane peptide were first combined to enable host–guest complexation. NorHA, DTT, lithium phenyl-2,4,6-trimethylbenzoylphosphinate (LAP) photoinitiator (4 mM final concentration), and PBS were subsequently added to the CDHA/adamantane mixture and vortexed thoroughly. Precursor solutions were pipetted into 8 mm wells of a silicone mold atop a glass slide, then photocrosslinked under 365 nm UV light (5 mW/cm^2^) for 5 min using a VWR UV crosslinker. After fabrication, hydrogels were swelled in PBS overnight at 37°C. The next day, hydrogels were mechanically characterized using the same rheology setup and tests as described above for lung characterization.

### 2.9 Statistics

All statistical analyses were performed using GraphPad Prism. Student’s t tests were utilized when comparing two experimental groups, while two-way ANOVA with Tukey’s HSD post hoc analysis was used for comparisons between more than two groups.

## 3. RESULTS

### 3.1 Aged mice exhibit increased tissue deposition with bleomycin administration compared to normal aged-matched controls

To investigate changes in lung tissue mechanics with development of pulmonary fibrosis, we utilized the well-established bleomycin model in aged mice [20,21,32]. 15-month-old C57BL/6J mice were trans-orally administered bleomycin (0.3 U/kg) and followed for 21 days before mechanically characterizing their lung tissue via rheology and nanoindentation (**Fig. 1**). Fibrosis development was evaluated through histopathology, where H&E staining was used to visualize cell and tissue accumulation in alveolar interstitial space, while Masson’s trichrome staining enabled analysis of collagen deposition (**Fig. 2**). We found that lungs from mice that received bleomycin exhibited increased overall tissue deposition and collagen content compared to control mice (Fig. 2A, B). The degree of fibrosis was quantified using trichrome-stained samples to determine Modified Ashcroft scores (Fig. 2C). Age-matched control lungs exhibited an average Ashcroft score of 3.3 +/- 1.07, compared to a significant increase of 4.5 +/- 0.97 in bleomycintreated lungs. Additionally, only 10% of fields of view in control mice had an Ashcroft score greater than 5, while this proportion increased more than threefold to 37% in the bleomycin-treated group. Furthermore, we performed immunohistochemical staining of lung tissue for type I collagen and hyaluronan binding protein (HABP). We observed increases in both ECM components in bleomycin-treated mouse lungs compared to control mice, consistent with previous studies (**Fig. S4**) [33–35]. Taken together, these results confirm the development of pulmonary fibrosis in bleomycin-treated aged mice.

**Figure 1.**
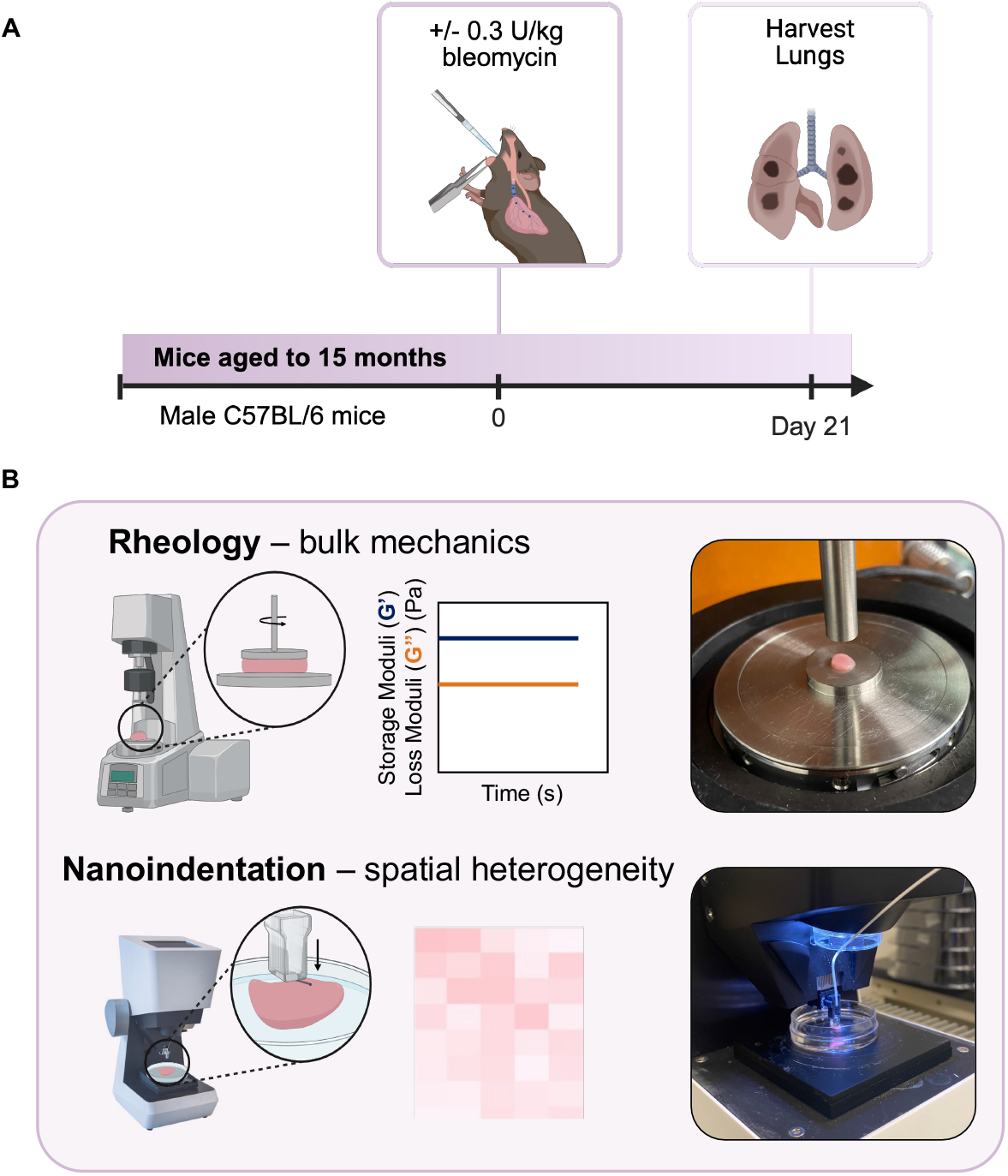
A) Bleomycin administration timeline. Male C57BL/6 mice were aged to 15 months before being orotracheally administered 0.3 U/kg bleomycin sulfate. After 21 days, mice were sacrificed and lungs were immediately processed for mechanical characterization. B) Left lung lobes were separated and utilized to evaluate bulk lung mechanics via rheology (top) and spatial mechanical heterogeneity via nanoindentation (bottom).

**Figure 2.**
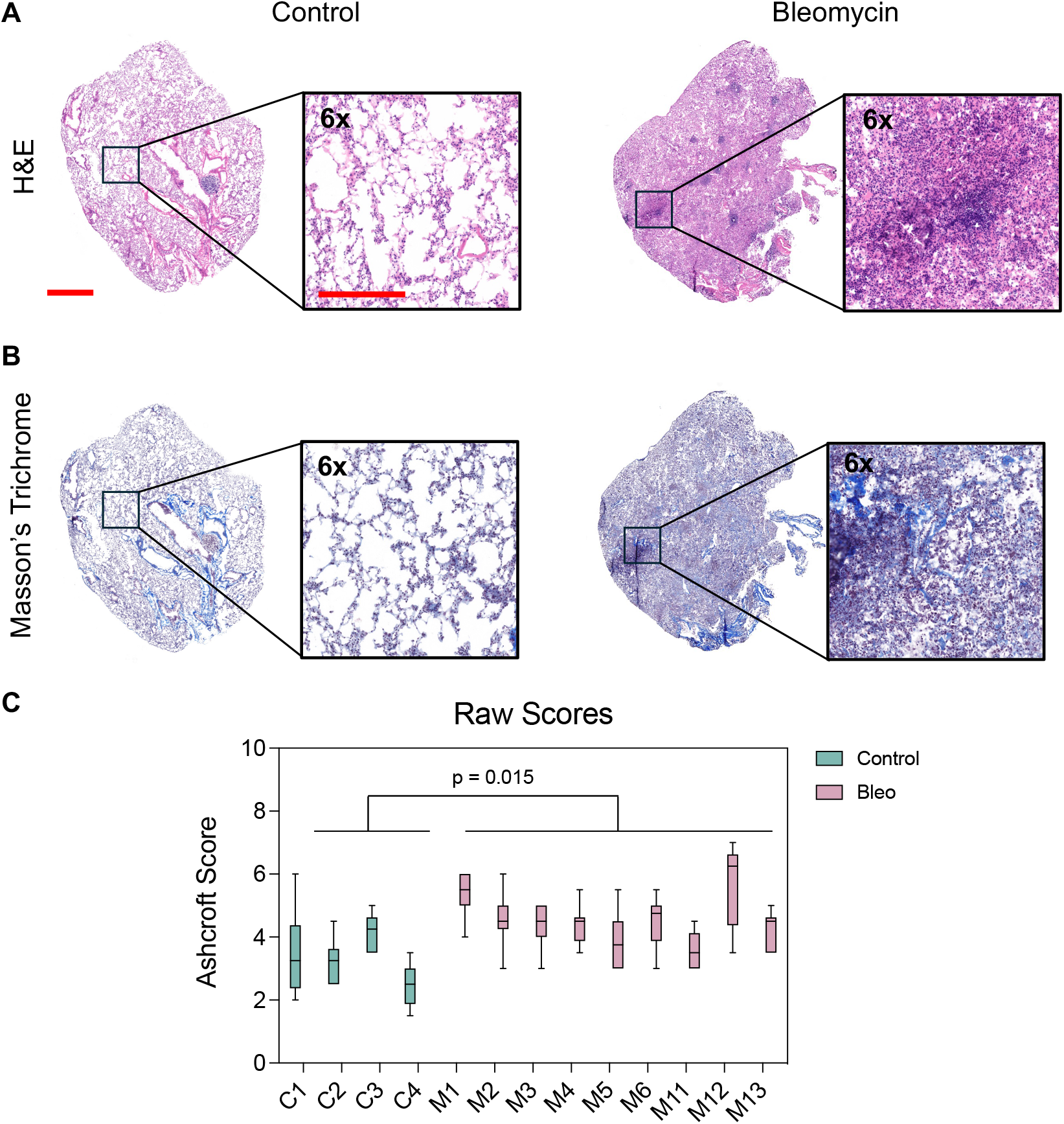
Histological evaluation of control and bleomycin-treated mouse lungs. A) Hematoxylin and eosin (H&E) staining for control and bleomycin-treated mouse lungs. B) Masson’s trichrome staining for control and bleomycin-treated mouse lungs. Scale bar = 1 mm; Inset scale bar = 50 μm. Extent of fibrosis was quantified by evaluating Masson’s trichrome-stained slides. C) Modified Ashcroft scores were assessed by two independently trained researchers who were double-blinded to image identities. Each boxplot represents the distribution of scores from 10 randomly selected 1 mm^2^ fields within a whole-slide image for each mouse. Whiskers indicate the minimum and maximum scores observed. Student’s t tests were used to assess differences in panel C.

### 3.2 Characterization of bulk lung tissue mechanics via rheology reveals increased lung stiffness but no change in lung viscoelastic behavior with bleomycin-induced fibrosis

We utilized oscillatory shear rheology to probe differences in bulk tissue mechanics between normal and fibrotic mouse lungs (**Fig. 3**). Time sweep rheological measurements were performed to assess the storage (G’) and loss (G”) moduli of lung tissue, granting insight into both tissue stiffness (G’) and viscoelastic behavior (G”/G’, tan delta). Bleomycin treatment resulted in varied bulk mechanical properties, with a general trend toward increased stiffness shown by an average Young’s modulus of about 4.7 kPa compared to control lungs at 2 kPa (Fig. 3A). We quantified no difference between tan delta values for control and bleomycin-treated groups, indicating that the viscoelastic character of lung was not changing with fibrosis. To further evaluate this, we tested the frequency-dependent mechanical behavior of control and bleomycin-treated mouse lungs, and again saw no significant difference between these groups, indicating that dynamic mechanical response was also not altered with fibrotic remodeling (Fig. 3B). We also investigated the stress relaxation behavior of lungs and found no difference in the time it took for lungs to relax half of the stress at a constant applied strain (τ_1/2_), providing further support that bleomycin-induced fibrosis did not significantly change lung viscoelasticity (Fig. 3C, D). While the heterogeneous, bimodal response seen in the bleomycin-treated group contributes to a wide range in stiffnesses and frequency-dependent responses, tan delta and τ_1/2_ values are notably consistent, with no trend observed between storage modulus (G’) and either tan delta (R^2^ = 0.03, p = 0.54) or τ_1/2_ (R^2^ = 0.21, p = 0.11) regardless of treatment group (Fig. 3E, F). Overall, these rheological measurements indicate that while lung stiffness increases with bleomycin-induced fibrosis, the viscoelastic behavior of tissue is not changing.

**Figure 3.**
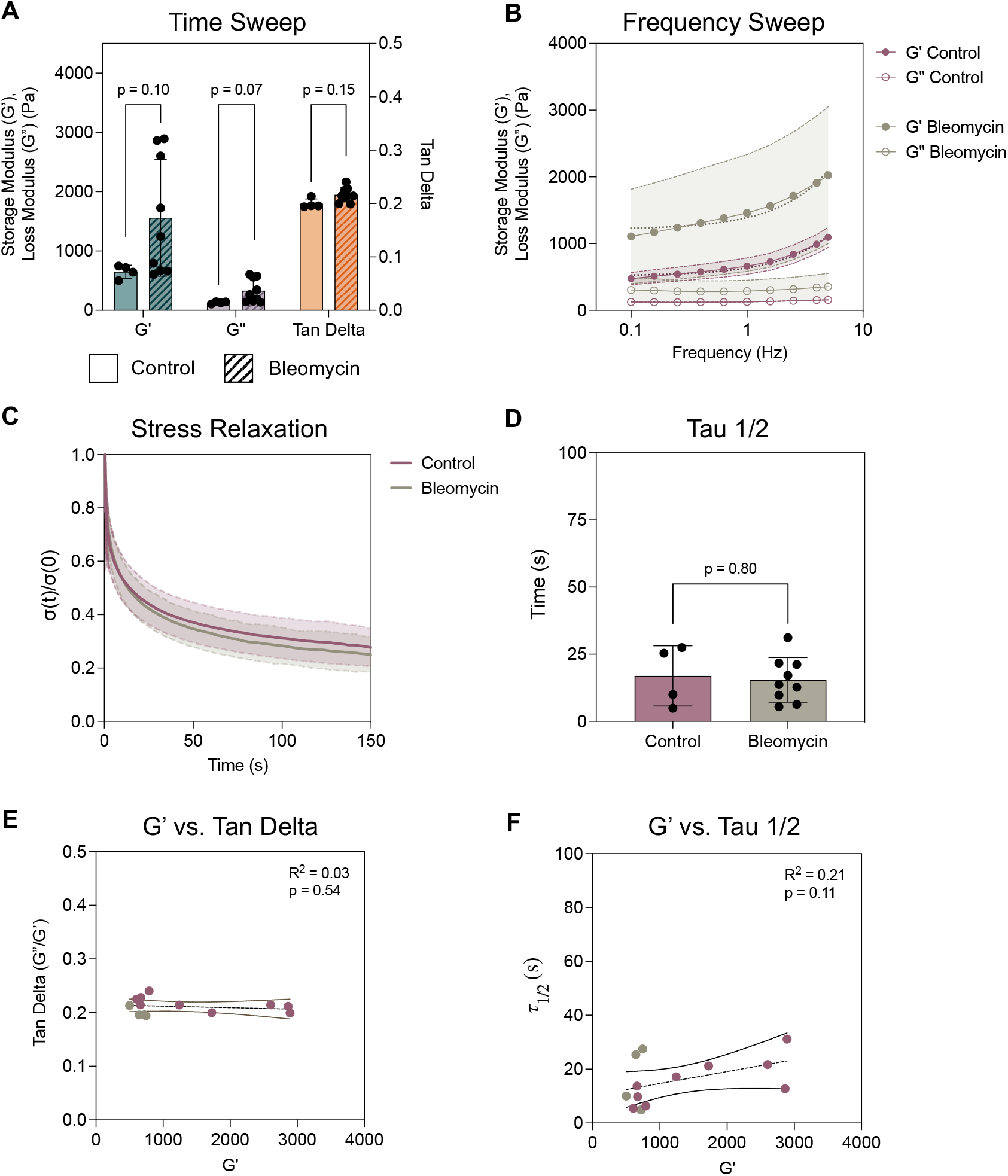
Rheological characterization of mouse lungs. A) Measurements of storage modulus (G’), loss modulus (G”), and tan delta (G”/G’) from a 30-second oscillatory time sweep reveal that while bleomycin treatment increases tissue stiffness (G’), viscoelasticity (tan delta) remains unaffected. B) The frequency-dependent mechanical response observed in both control and bleomycin-treated lungs further supports the preservation of viscoelastic behavior across a range of frequencies from 0.1-10 Hz. C) Control and bleomycin-treated mouse lungs display similar extent of stress relaxation (stress over time, σ(t), normalized to initial stress, σ(0)) and D) time to relax half of the applied stress (τ_1/2_), revealing that regardless of bleomycin treatment, viscoelasticity is maintained. E) Evaluation of stiffness (G’) versus tan delta and F) stiffness versus τ_1/2_ reveals no evident correlation between measurements, highlighting the consistency of viscoelasticity in normal and fibrotic lung. Shaded regions represent standard deviation for datasets. Each point represents one lung average. Student’s t tests were used to assess differences between groups.

### 3.3 Nanoindentation highlights spatial heterogeneity in bleomycin-treated mouse lungs and further supports no change in lung viscoelasticity

To further spatially resolve mechanical properties of normal and fibrotic lung tissue, we employed nanoindentation to perform similar analyses as those done with rheology (**Fig. 4**). For each lung, storage and loss moduli were measured at 64 points across an 8 x 8 matrix with a 200 µm step size, covering approximately 1.1 mm^2^ of tissue per sample. These point-based measurements were used to calculate an average stiffness for each lung, where we observed a significant increase in lung stiffness with bleomycin treatment compared to controls (Fig. 4A). While we observe significantly higher average storage moduli (E’) with nanoindentation (Fig. 4A) than with rheological characterization (Fig. 3A), this is likely a consequence of differences in measurement modes and scale. Rheology captures bulk tissue mechanics, averaging across both solid tissue and alveolar airspace, where nanoindentation is inherently biased toward solid regions, as measurements require direct probe contact with the tissue surface. Utilizing this approach enables us to characterize local mechanical changes within a lung, providing a more detailed map of heterogeneity and complementing bulk measurements to better understand how fibrosis alters tissue mechanics.

**Figure 4.**
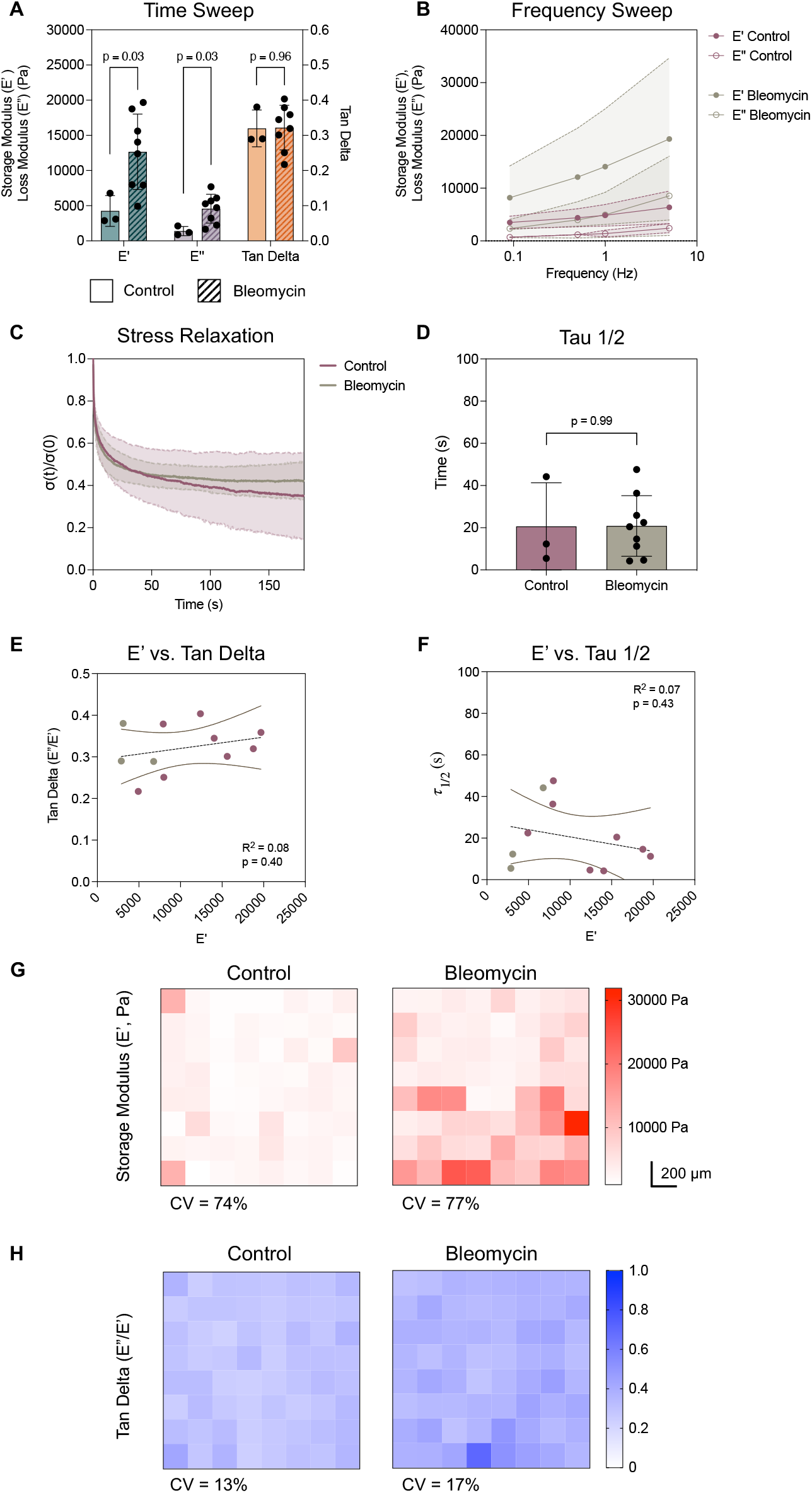
Nanoindentation evaluation of mouse lung mechanics. A) Average storage moduli (E’), loss moduli (E”), and tan delta (E”/E’) from an 8 x 8 matrix scan across each lung measured at 1 Hz. Each point represents the average stiffness per lung. Bleomycin-treated mouse lungs exhibit a significant increase in stiffness (E’) while maintaining similar viscoelastic behavior (tan delta) to control lungs. B) Control and bleomycin-treated mouse lungs exhibit frequency-dependent mechanics across a range of 0.1-10 Hz, demonstrating viscoelastic behavior. C) Control and bleomycin-treated mice exhibit comparable levels of stress relaxation (stress over time, σ(t), normalized to initial stress, σ(0)), and D) the time required to reach half of the applied stress (τ_1/2_), demonstrating that despite bleomycin treatment, lung tissue viscoelasticity is preserved. Shaded regions represent standard deviation for datasets. Each point represents one lung average. Statistical analyses performed via Student’s t tests. ∗*p* < 0.05. E) Evaluation of stiffness (E’) versus tan delta and F) stiffness versus τ_1/2_ reveals no evident correlation between measurements, highlighting the consistency of viscoelasticity in normal and fibrotic lung. G) Representative mechanical maps of storage moduli (E’) from control and bleomycin-treated mouse lungs demonstrate high levels of heterogeneity and are quantified by coefficients of variation for each group (CV, average / standard deviation). H) Representative mechanical maps of tan delta show consistent measurements within samples, which is quantitatively shown through lower CVs than stiffness maps. Each voxel represents a 50 µm indentation measurement area, with a 200 µm step size between points.

Again, we saw no difference in tan delta values between control and bleomycin-treated groups, indicating no change in lung viscoelasticity (Fig. 4A). To further assess viscoelastic differences between normal and bleomycin-treated mouse lungs, we conducted frequency sweep and stress relaxation tests. These measurements complemented our bulk rheological analysis and revealed no significant changes in frequency-dependent or stress relaxation behavior between groups (Fig. 4B–D). Furthermore, while we again observe a heterogeneous distribution in bleomycin-treated mouse lung stiffnesses, there is still no significant difference in tan delta or τ_1/2_ values, nor is there a correlation between storage modulus (E’) and these measurements (Fig. 4E, F). Heatmaps of nanoindentation measurements demonstrated high levels of heterogeneity in storage modulus within samples (Fig. 4E). In contrast, tan delta values were more uniform across samples, indicating consistent viscoelastic behavior in both normal and bleomycin-treated mouse lungs (Fig. 4F). This distinction was evident by quantitative analysis of variation, as storage modulus values exhibited significantly higher coefficients of variation (CV ∼ 75%) compared to tan delta (CV ∼ 15%), highlighting greater heterogeneity in stiffness than in viscoelasticity (Fig. 4E, F). Taken together, these findings support the conclusion that lung viscoelasticity is not altered by bleomycin-induced fibrotic remodeling.

To determine whether the preservation of viscoelasticity in fibrotic lungs was specific to aged mice, we extended our nanoindentation analysis to young (12-week-old) normal and bleomycin-treated mice (**Fig. S5**). Similar to aged animals, young mice exhibited significantly increased lung stiffness following bleomycin treatment (Fig. S5A, S5B). Despite this increase in stiffness, there were no significant changes in tan delta (E″/E′), indicating that viscoelastic behavior remained consistent (Fig. S5A, S5C). Moreover, heterogeneity within young mouse samples followed similar trends to those observed in aged mice, with higher coefficients of variation within stiffness measurements for each lung, while tan delta variability was consistent with aged mice (Fig. S5D, S5E). Together, these findings suggest that fibrotic stiffening occurs without significantly altering viscoelastic behavior, regardless of age.

### 3.4 Hyaluronic acid-based hydrogels recapitulate normal and fibrotic lung mechanics

Adapting a tunable HA-based hydrogel system previously developed in our lab [8,23,36], we formulated hydrogels that mimicked the mechanics of control and bleomycin-treated mouse lungs (**Fig. 5**). By tuning polymer weight percent, crosslinker concentration, and incorporating physical guest-host interactions to impart viscoelasticity, we engineered hydrogels with storage (G’) and loss (G”) moduli equivalent to those of lungs from control and bleomycin-treated mice, as measured by rheology (Fig. 5A, B). In addition to matching lung stiffness, we recapitulated more complex mechanical cues like tissue viscoelasticity with this hydrogel system. This is demonstrated by comparable tan delta values between hydrogels and lungs (Fig. 5A, B), frequency-dependent mechanical behavior (Fig. 5C, D), as well as a similar extent of stress relaxation of about 80% (Fig. 5E, F). Taken together, these results demonstrate the unique ability of hydrogels to model static and dynamic tissue mechanics in the context of health and disease.

**Figure 5.**
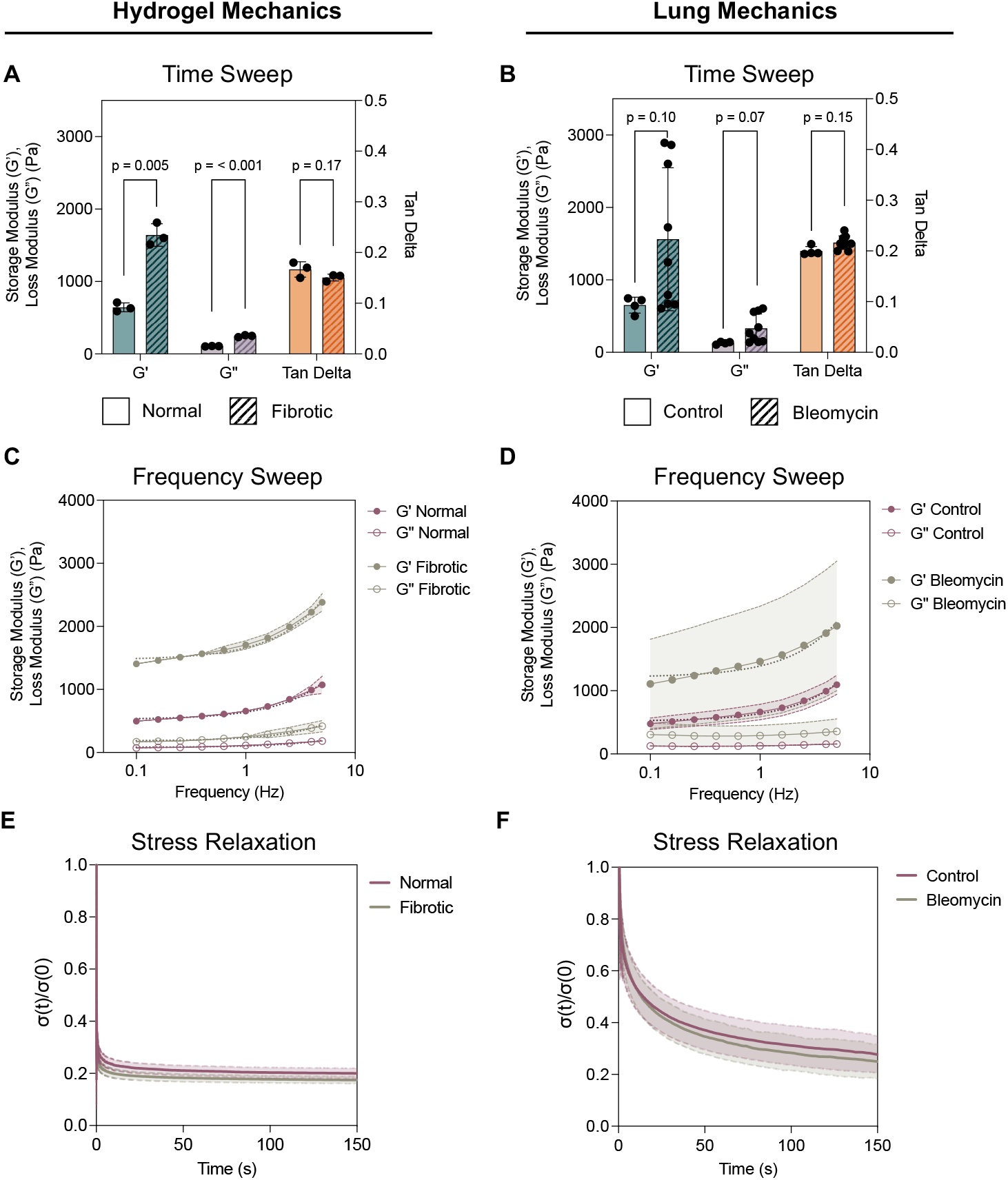
Rheological comparison of hyaluronic acid-based hydrogels with normal and bleomycin-treated mouse lungs. Measurements of storage modulus (G’), loss modulus (G”), and tan delta (G”/G’) from a 30-second oscillatory time sweep demonstrate the capacity of A) hydrogel systems to mimic stiffness and viscoelastic behavior of B) normal and fibrotic lungs. Frequency sweeps (0.1-10 Hz) show similar frequency-dependent behavior between C) hydrogels and D) lung tissue. E) Hydrogels exhibit similar extent of stress relaxation (stress over time, σ(t), normalized to initial stress, σ(0)) to F) normal and fibrotic mouse lungs during 3 min of a constant 5% applied strain. Lung mechanical data are reproduced from Fig. 2. N = 3 hydrogels per group.

## 4. DISCUSSION

Our findings are consistent with previous studies reporting heterogeneous fibrotic responses to bleomycin administration in mice, a well-documented feature of this model that reflects both the extent of fibrosis as well as the spatial distribution [15,37–39]. This heterogeneity may result from differences in local delivery of bleomycin, differences in immune response, or tissue susceptibility, among other considerations [15,37–39]. Although this heterogeneity led to a wide range of lung stiffness values within our datasets, the bimodal distribution observed in both our rheological and nanoindentation measurements parallels the distribution of Young’s moduli reported in IPF patient lungs [40]. Thus, while the heterogeneity observed in extent of fibrosis is a disadvantage of this model, it also recapitulates the non-uniform progression of fibrotic remodeling in this disease. Strikingly, while storage modulus measurements were highly variable, our primary measures of viscoelasticity were remarkably consistent between both young and aged control and bleomycin-treated mice (**Figs. 3, 4, S5**).

These findings suggest that, despite ECM remodeling and collagen accumulation in bleomycin-treated lungs (**Fig. S4**), lung tissue retains its ability to dissipate mechanical energy and recover from deformation (**Figs. 3, 4**). This preservation of viscoelastic behavior may reflect compensatory changes in ECM composition—particularly the overdeposition of uncrosslinked components such as hyaluronic acid, which has been shown to increase in fibrotic lungs [33,41,42]. Considering these findings, our results support the potential of HA as a contributor to preserved viscoelastic tissue mechanics in fibrosis. Given that viscoelasticity is essential for normal lung function, our results point to a potentially underrecognized feature of fibrotic remodeling. Furthermore, while prior studies have reported Young’s moduli and some dynamic mechanical properties (resistance, elastance, hysteresivity) for normal and fibrotic lungs, to our knowledge, this study is the first to compare the mechanical properties of normal and fibrotic lungs from the bleomycin model of pulmonary fibrosis using both bulk and spatially resolved characterization techniques [22,43,44]. These insights could be leveraged to inform the design of fibrosis therapeutics aimed not only at reducing tissue stiffness but also at preserving the dynamic mechanical properties of lung tissue. Additionally, we demonstrated that hydrogel systems can effectively recapitulate these mechanical properties. We previously developed high-throughput methods to engineer hydrogels, providing a more physiologically-relevant platform for therapeutic testing compared to conventional tissue culture plastic or glass [31].

Our hydrogel models recapitulated the storage (G’) and loss (G”) moduli, tan delta (G”/G’), and extent of stress relaxation of normal and bleomycin-treated mouse lungs. Utilizing these systems could enable experimental interrogation of how cell behavior changes throughout fibrotic diseases progression in a controlled, mechanically relevant environment. However, our engineered hydrogels exhibited faster relaxation timescales than tissue (**Fig. 5**). Several studies have investigated this behavior in hydrogels by incorporating polymers of varying molecular weights, utilizing fibrillar collagen, or introducing reversible bonds to modulate the timescales of stress relaxation [45–48]. These studies demonstrate that cells cultured on substrates with different relaxation timescales exhibit different morphologies, proliferation rates, remodeling of surrounding matrix, and even different levels of osteogenic differentiation in the case of mesenchymal stem cells [45–48]. Considering these results, though the hydrogel system we employed here exhibits faster relaxation than tissue, we could utilize similar methods to incorporate different relaxation timescales in the future. Additionally, while we demonstrated hydrogels can capture the bulk mechanical properties of normal and fibrotic lungs measured by rheology, we did not attempt to recreate the heterogeneity observed at the microscale with nanoindentation. To incorporate this property into our hydrogel system, we could leverage photopatterning. Our lab and others have previously utilized this technique to fabricate hydrogels that exhibit different stiffnesses and levels of viscoelasticity within the same hydrogel [8,49,50]. Taken together, adjusting hydrogel stress relaxation timescales and/or spatial heterogeneity could be incorporated into future iterations of this hydrogel system to further model complex tissue mechanics.

While this study employed an aged mouse model, as prior research showed it more accurately reflected the chronic and progressive nature of pulmonary fibrosis in humans [20,21], we also assessed whether mechanical changes were conserved in young mice. Our findings indicate that bleomycin-induced fibrotic stiffening occurs in both young and aged lungs, yet viscoelastic properties, as measured by tan delta, remain unchanged (**Figs. 3, 4, S5**). This preservation of viscoelasticity, despite increased stiffness, suggests a potentially fundamental feature of fibrotic lung mechanics. Notably, while both young and aged mice exhibited similar intra-lung heterogeneity in stiffness, viscoelasticity measurements were consistent, reinforcing the robustness of this preserved mechanical behavior. To our knowledge, this is the first study to demonstrate that viscoelastic tissue behavior is maintained in fibrotic mouse lungs independent of age, underscoring the importance of considering both elastic and time-dependent mechanical properties when modeling or evaluating lung fibrosis.

## 5. CONCLUSIONS

We utilize the bleomycin mouse model of fibrosis to investigate mechanical changes in normal and fibrotic lungs. By employing rheological and nanoindentation methods to characterize tissue, we are able evaluate differences in bulk and spatially-resolved tissue mechanics. We found that while tissue stiffness increased with fibrosis progression, viscoelastic behavior was preserved. This finding was upheld in both rheological and nanoindentation measurements, and across young and aged mouse models. Furthermore, while both rheology and nanoindentation underscored the heterogeneity in fibrotic tissue stiffness, viscoelasticity measurements were remarkably consistent. Using the same rheological tests that we used to characterize mouse lungs, we demonstrated the ability of hyaluronic acid-based hydrogels to recapitulate static and dynamic mechanical properties of tissue. Overall, these findings reveal that dynamic tissue mechanical properties such as viscoelasticity do not always align with static measures like stiffness, emphasizing the critical need for *in vitro* models that robustly mimic complex aspects of tissue mechanics to enable the development of more effective therapeutics.

## 6. ACKNOWLEDGEMENTS

We thank Dr. Jenna Sumey for her contributions to animal care, mechanical characterization of lung tissue, and support in this work, as well as the lab of Dr. Christopher Highley for allowing use of their rheometer to perform lung and hydrogel mechanical characterization. We thank the UVA Research Histology Core for cryopreservation, sectioning, and H&E and Masson’s trichrome staining of lung samples, and the UVA Advanced Microscopy Facility for imaging support. Both cores are supported by the University of Virginia School of Medicine, with Research Resource Identifiers (RRIDs): SCR_025470 and SCR_01873, respectively. This work was supported by the NIH (R35GM138187 to S.R.C, R01HL179312 to J.M.S., and F32HL170760 to R.T.H.) and NSF (GRFP to L.R.A. and M.L.S.). The content is solely the responsibility of the authors and does not necessarily represent the official views of the National Institutes of Health.

