## Supplemental Figures for "Lung tissue viscoelasticity is preserved with bleomycin-induced fibrosis in mice"

### Supplemental Information

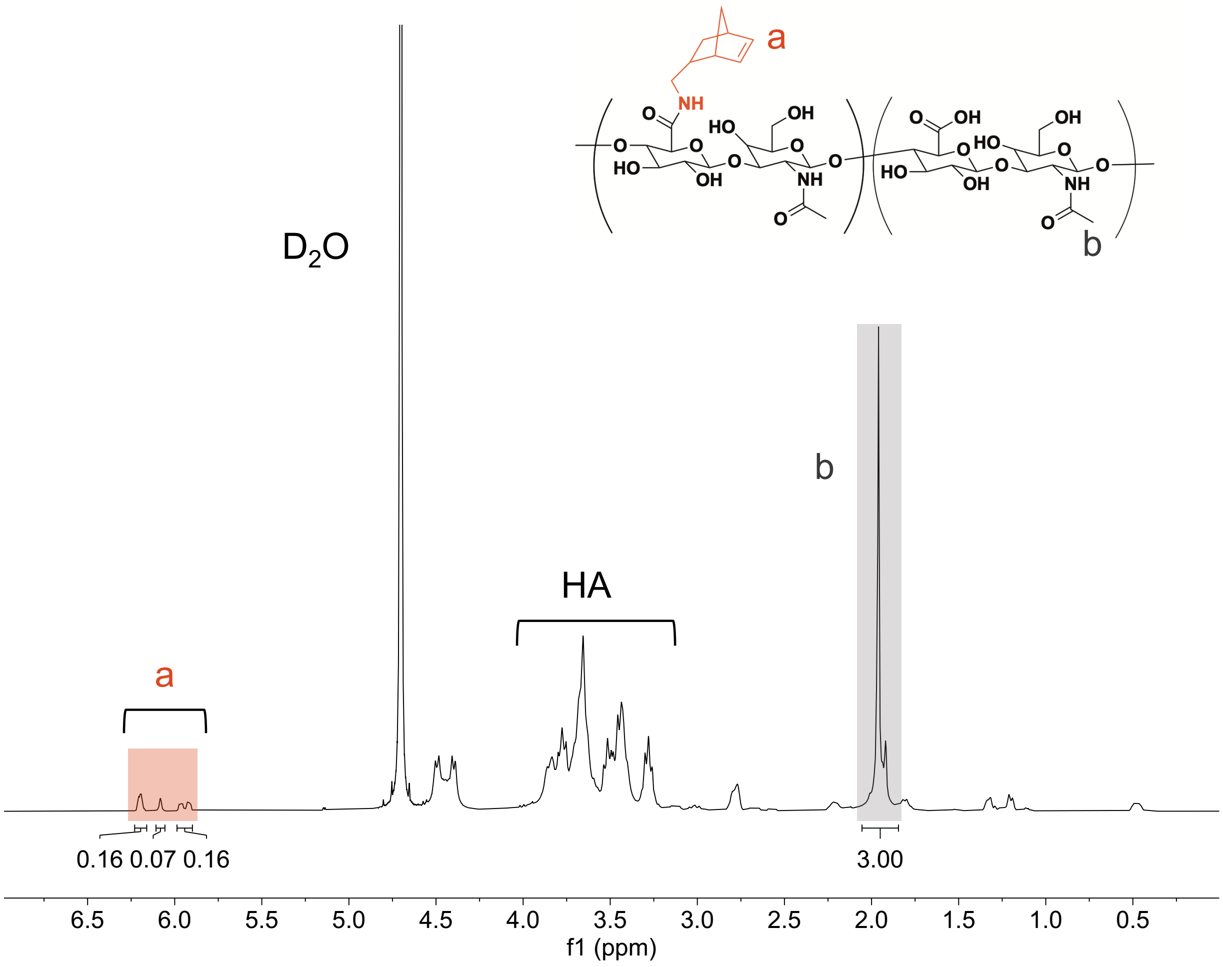


**Figure S1.** $^1$H NMR spectra of norbornene-modified hyaluronic acid (NorHA). Modification of HA with pendant norbornenes was determined to be 19.5%, indicated by the integration of peaks at $\delta$ = 5.8, 6.05, and 6.2 ppm (2H, ‘a’) normalized to the *N*-acetyl group on HA (3H, ‘b’).

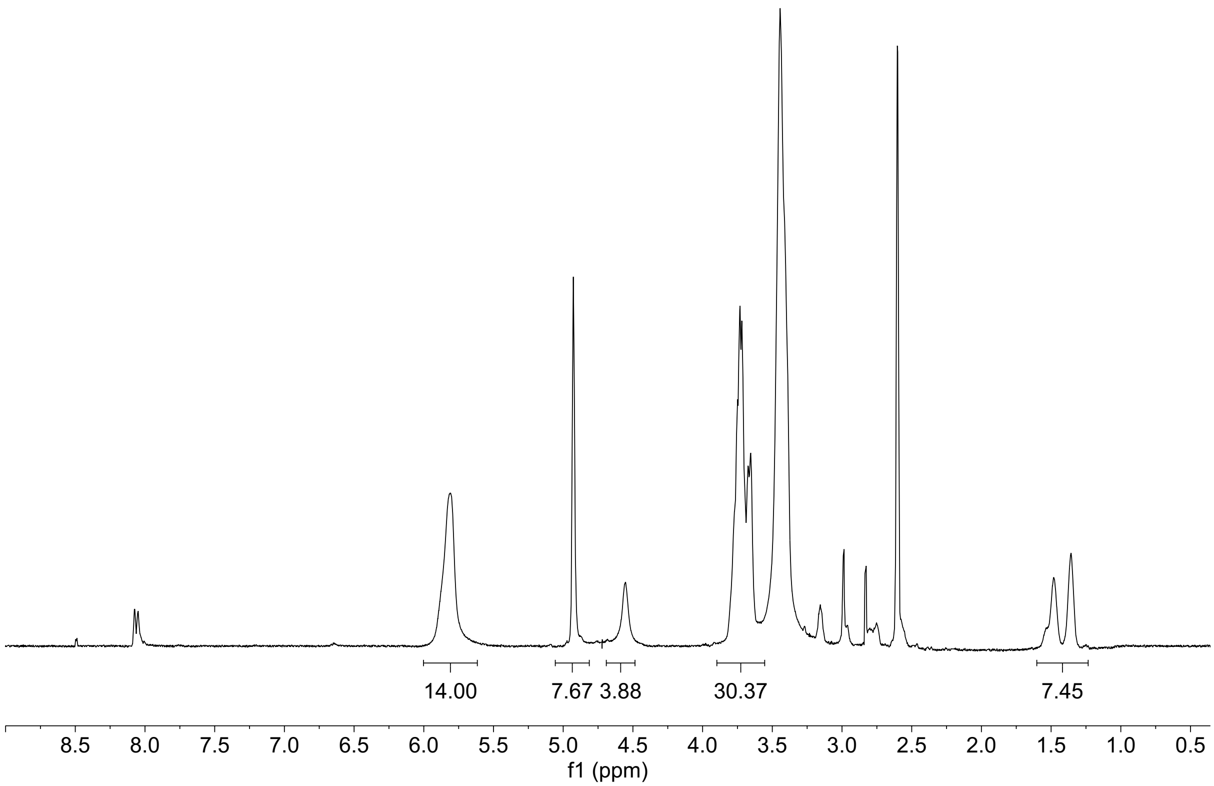


**Figure S2.** $^1$H NMR spectra of 6-(6-aminohexyl)amino-6-deoxy-β-cyclodextrin (CD-HDA). Modification of β-CD with HDA was determined to be 62%, indicated by the integration of peaks at $\delta$ = 1.14-1.6 ppm (12H).

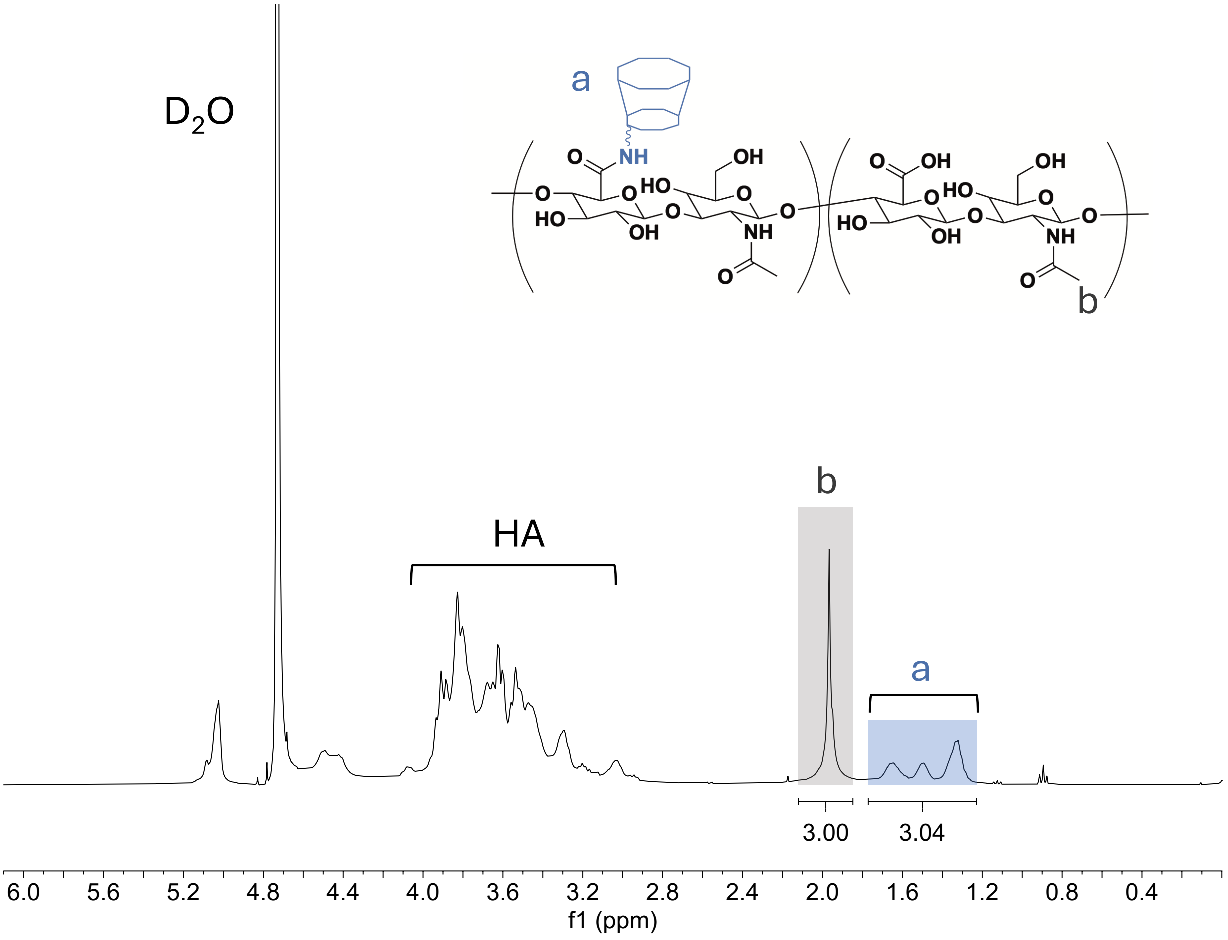


**Figure S3.** $^{1}$H NMR spectra of β-cyclodextrin-modified hyaluronic acid (CDHA). Modification of HA with pendant cyclodextrins was determined to be about 25% by integration of hexane linker peaks at $\delta$ = 1.2-1.7 ppm (12H, a) relative to the *N*-acetyl group of HA (3H, b).

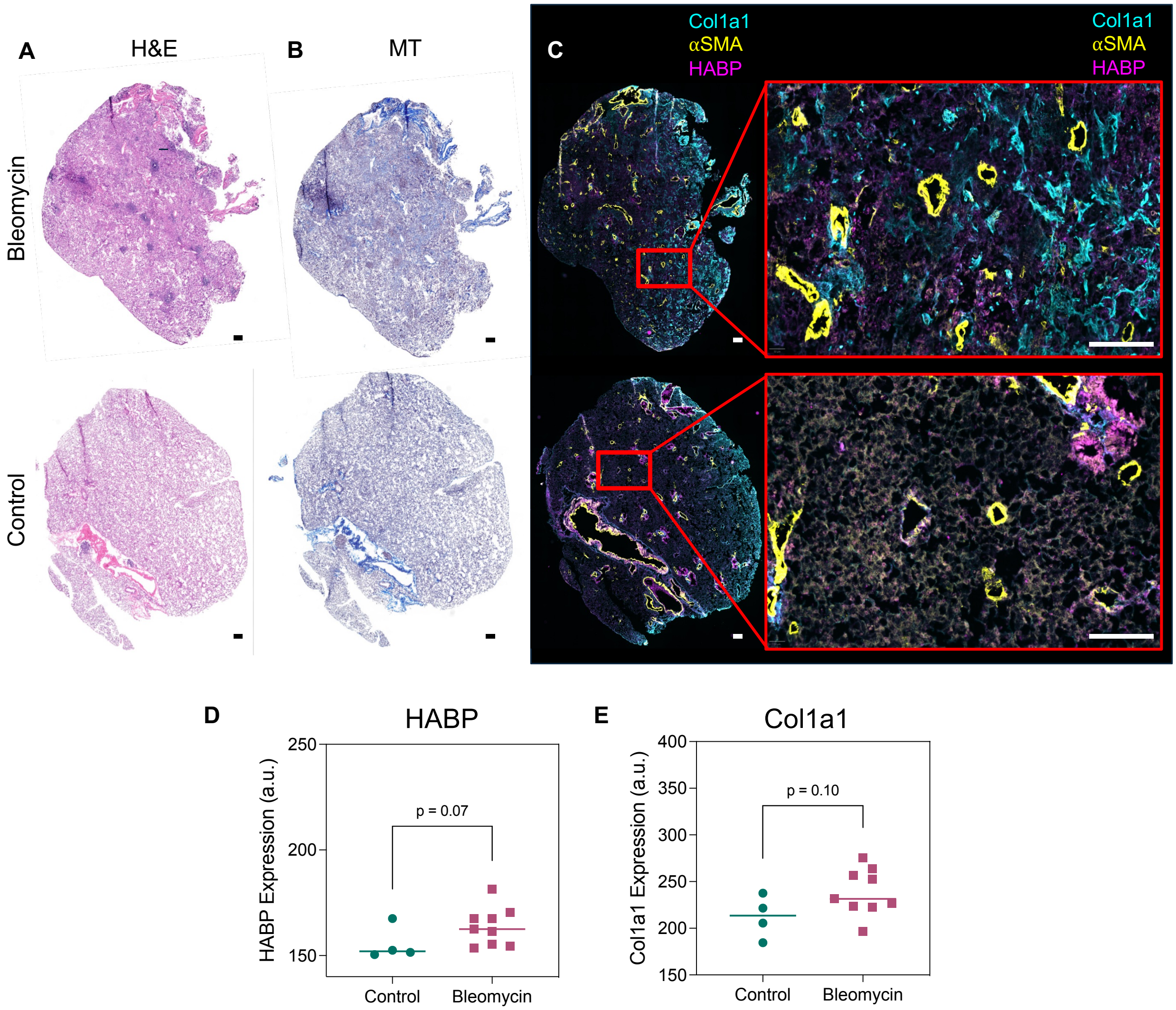


**Figure S4.** Characterization of extracellular matrix composition in control and bleomycin-treated mouse lungs. A) Hematoxylin and eosin (H&E) and B) Masson's trichrome staining for control and bleomycin-treated mouse lungs. C) Immunofluorescent staining shows a relative increase in type 1 collagen (Col1a1, cyan), alpha smooth muscle actin (αSMA, yellow), and hyaluronan binding protein (HABP, magenta) in bleomycin-treated mouse lungs compared to controls. All scale bars = 200 μm. Quantification of D) HABP and E) Col1a1 in control and bleomycin-treated mouse lungs demonstrate an increasing trend in protein levels in fibrotic lung. Student's t tests were used to compare groups. Each dot represents one mouse with lines indicating the median for each group.

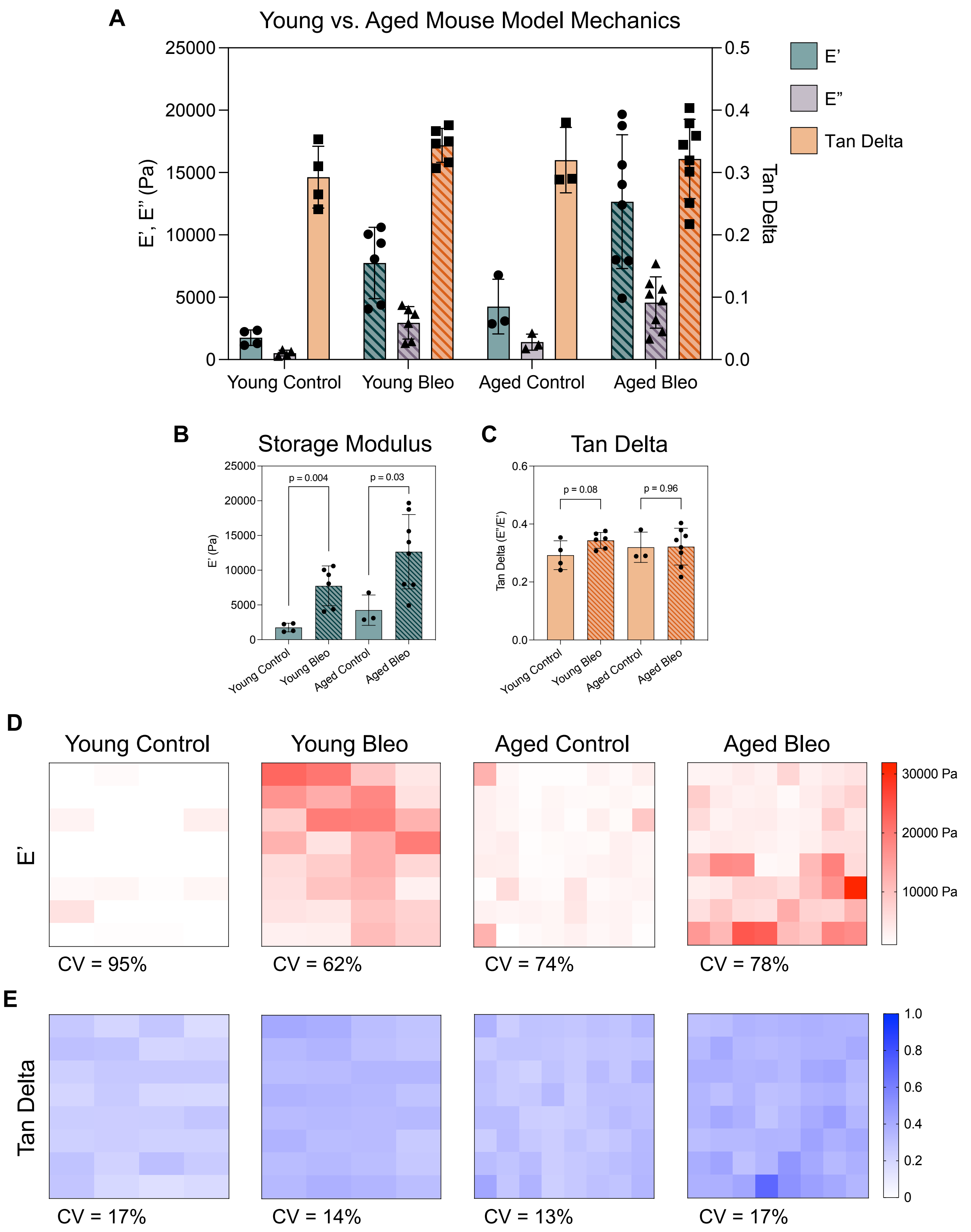


**Figure S5.** Comparison of young (12 week-old) and aged (15 month-old) normal and bleomycin-treated lung mechanics. Aged data reproduced from Figure 4. A) Average storage moduli (E'), loss moduli (E"), and tan delta (E"/E') from a matrix scan across each lung measured at 1 Hz. Each point represents the average stiffness per lung. B) Both young and aged mouse lungs exhibit a significant increase in storage modulus with bleomycin treatment. C) There are no significant differences in tan delta regardless of mouse age or bleomycin treatment, highlighting the persistence of viscoelasticity. Each point represents one lung average. Statistical analyses performed via Student's t tests. D) Representative mechanical maps of storage moduli (E') from young and aged control and bleomycin-treated mouse lungs demonstrate high levels of heterogeneity and are quantified by coefficients of variation for each group (CV, average / standard deviation). F) Representative mechanical maps of tan delta show consistent measurements within samples, which is quantitatively shown through significantly lower CVs than stiffness maps. Each voxel represents a 50 μm indentation measurement area, with a 200 μm step size between points.
